# Discrimination Exposure and DNA Methylation of Stress-Related Genes in Latina Mothers

**DOI:** 10.1101/306027

**Authors:** Hudson P Santos, Benjamin C Nephew, Arjun Bhattacharya, Xianming Tan, Laura Smith, Reema Alyamani, Elizabeth M Martin, Krista Perreira, Rebecca C Fry, Christopher Murgatroyd

## Abstract

Latina mothers, who have the highest fertility rate among all ethnic groups in the US, are often exposed to discrimination. The biological impacts of this discrimination are unknown. This study is the first to explore the relationship between discrimination and DNA methylation of stress regulatory genes in Latinas. Our sample was Latina women (n = 147) with a mean age of 27.6 years who were assessed at 24-32 weeks’ gestation (T1) and 4-6 weeks postpartum (T2) and reside in the U.S. Blood was collected at T1, and the Everyday Discrimination Scale (EDS) was administered at T1 and T2. DNA Methylation at candidate gene regions was determined by bisulphite pyrosequencing. Associations between EDS and DNA methylation were assessed via zero-inflated Poisson models, adjusting for covariates and multiple-test comparisons. Discrimination was negatively associated with methylation at CpG sites within the glucocorticoid receptor (*NR3C1*) and brain-derived neurotrophic factor (*BDNF*) genes that were consistent over time. In addition, discrimination was negatively associated with methylation of a CpG in the glucocorticoid binding protein (*FKBP5*) at T1 but not at T2. This study underscores the complex biological pathways between discrimination and epigenetic modification in Latina women that warrant further investigation to better understand the genetic and psychopathological impact of discrimination on Latino mothers and their families.

## 1. Introduction

The accumulation of stress over the lifespan can contribute to biological vulnerability and directly affect health outcomes for mothers and their children. Latina women, who have the highest fertility rate among all ethnic groups and represent the largest minority group in the US (Center, 2015), are exposed to a multitude of stressful events and sociocultural factors, including discrimination (Ayón, 2015). Extant research in Latinas has largely focused on varied levels of exposure to risk and protective factors in the perinatal period including socio-determinants of health (e.g., socioeconomic background), prenatal care, social support, and stress. These factors, however, do not adequately account for all of the noted disparities in perinatal outcomes, such as perinatal depression and morbidity (Guintivano et al., 2017; Howell et al., 2017).

Discrimination has been defined as differential treatment based on: (1) race that disadvantages a racial/ethnic group and/or (2) inadequately justified factors other than race/ethnicity that disadvantages a racial/ethnic group (Council, 2004). A contributing factor in health disparities and social inequality, discrimination has been associated with several adverse physical and mental health outcomes in minority groups (Wallace et al., 2016; Williams and Mohammed, 2009). A recent meta-analysis of 150 studies demonstrated a statistically significant effect size of racial discrimination on health, with the largest effect on mental health (r = .20, 95% CI: .17, .24) (Carter et al., 2017). Earlier meta-analyses found similar associations (Lee and Ahn, 2012), and discrimination is a significant predictor of poor mental health in Black and Latino immigrants (Gee et al., 2006). A UK study concluded that cumulative exposure to racial discrimination has incremental long term effects, and noted that assessing discrimination at only one time point may underestimate the adverse effects of discrimination on mental health (Wallace et al., 2016). Among Latinos, discriminatory experiences are specifically associated with decreased self-esteem and emotional stress, increased anxiety and depressive symptoms, and social isolation (Ayón, 2015). A recent nationally representative survey suggested that one in three Latinos report discrimination based on ethnicity, and one in five report that they have avoided seeking medical care or calling police authorities because they were concerned that they or a family member would experience discrimination (Health et al., 2017). Overall, these studies suggest that Latinos face discrimination across the US. Moreover, in the current political climate, discrimination against Latinos may be increasing (Almeida et al., 2016).

Several mechanisms for the adverse effects of discrimination on mental health have been described (Berger and Sarnyai, 2015). This study focuses on discrimination as a potent stressor that can result in neuroendocrine dysregulation, via DNA methylation, of stress regulatory genes which can ultimately have deleterious health effects. DNA methylation is the addition of a methyl group, usually to cytosines within CpG dinucleotides, which when located in promoter regions generally represses gene expression. Stress reactivity has been hypothesized to mediate the impact of the social environment on health. Exposure to various environmental stressors can alter the expression of hypothalamic–pituitary– adrenal axis (HPA) associated genes, and social adversity has potent effects on stress-regulating pathways, resulting in altered stress responsivity. Dysregulation of the stress response, notably expression of HPA genes through epigenetic modifications such as DNA methylation, is likely to contribute to stress-related health disparities and provide a link between the stressful social environment and disease development (Mitchell et al., 2016; Szyf, 2013).

Little is known about how perceived racial discrimination relates to DNA methylation patterning. A recent study reported an inverse relationship between perceived racial discrimination and DNA methylation at seven CpG sites (six genes related to tumor suppression protein-coding) in African-American women enrolled in a blood pressure study (Mendoza et al., 2018). To our knowledge, it is unknown if there are similar associations between discrimination and DNA methylation in Latinos. However, studies have reported dysregulation of the stress response in depressed Latina mothers (Lara-Cinisomo et al., 2017), supporting the rationale for the present investigation of discrimination and DNA methylation.

Two genes critically involved in the regulation of the stress response are the glucocorticoid receptor gene (*NR3C1*) (Herman et al., 2012) and the glucocorticoid receptor chaperone protein gene *FKBP5* (Zannas et al., 2015). DNA methylation at key specific CpG sites within *NR3C1* exon 1-F promoter and two key CpGs within intron 7 of *FKBP5* have been associated with early adversity (Palma-Gudiel et al., 2015). In addition, stress induced neuroplasticity associated with altered HPA function is mediated by functional interactions between glucocorticoids and brain-derived neurotrophic factor (*BDNF*) (Numakawa et al., 2017) where methylation at a key promoter region IV has been linked to environmental stressors in humans and rodent models (Mitchelmore and Gede, 2014). Based on this integrated literature, we postulated that one biological mechanism implicated with discrimination is neuroendocrine-associated dysregulation via epigenetic programming of stress regulatory genes. The current study investigated associations between methylation at specific CpG sites within the *NR3C1, FKBP5,* and *BDNF* genes and perceived discrimination during pregnancy and early postpartum in a population of Latina mothers. It was hypothesized that DNA methylation within the *NR3C1, FKBP5,* and *BDNF* genes is inversely associated with perceived discrimination in Latina women in the U.S.

## 2. Material and Methods

### 2.1 Participants

Healthy pregnant Latina women (n = 150) living in North Carolina (NC) were enrolled in the study between May 2016 to March 2017. Eligibility criteria included: (1) 18-45 years old, (2) Spanish- or English-speaking, (3) carrying a singleton pregnancy, (4) available for follow-up at 6 weeks postpartum. Exclusion criteria were: (1) currently experiencing severe depressive symptoms as determined by psychiatric interview, (2) history of psychotic or bipolar disorder, or receiving psychotropic therapy, (3) substance dependence in the last two years, (4) fetal anomaly, or (5) life-threatening conditions. These exclusions were adopted to avoid confounders and control for severe mood symptoms with onset before the study time frame. Data collection was completed in English or Spanish, depending on participants preference, by a trained research assistant at the prenatal visit at 24-32-week gestation (T1) and at 4-6 weeks postpartum (T2). All measures used in this study had validated versions in English and Spanish. The Institutional Review Board of the University of North Carolina at Chapel Hill approved this study (#15-3027).

### 2.2 Measures

#### 2.2.1 Perceived Discrimination

The Everyday Discrimination Scale (EDS), a nine-item questionnaire, was used to measure routine, day-to-day experiences of discrimination at T1 and T2. The stem question is: “In your day-to-day life, how often do any of the following things happen to you?” Sample items include: “You are treated with less courtesy than other people are,” “People act as if they think you are dishonest” and, “You are called names or insulted.” Participants then link main reason for these discrimination experiences, including gender, race, ancestry, religion. In addition, participants were asked whether they have felt any type of ethnicity-based discrimination during their lifetime. The EDS is a widely used measure of subjective experiences of discrimination (Williams et al., 1997), with validated Spanish translation (Campo-Arias et al., 2015; Park et al., 2018). It correlates with measures of institutional racial discrimination and interpersonal prejudice (Krieger et al., 2005) and does not prime the subjects to think about race, which limits cues to prejudice prior to responding to the questions (Deitch et al., 2003). The 9-item Likert response scale for frequencies ranged from 0 (“never”) to 5 (“almost every day”). We constructed a mean summary that ranged from 0 to 5, with a higher score indicating a higher frequency of perceived discrimination. Cronbach’s alpha for item consistency for the EDS in our sample was 0.86 for T1 and 0.89 for T2.

#### 2.2.2 DNA Methylation

To minimize variability in stress, the study blood draw was incorporated into the routine prenatal blood draw at T1 followed by self-report measures. A 6 ml blood sample was drawn from a peripheral vein into a chilled EDTA-vacutainer, placed immediately on ice and processed. The buffy coat was separated by centrifugation, frozen on dry ice, and stored at −80^°^C at the University of North Carolina Biobehavioral lab until DNA extraction. DNA extraction was performed with the QIAamp DNA Blood Mini Kit and extracted DNA was stored at −80^°^C in individual cryovials until shipment. The extracted DNA was transported on dry ice to the UK for DNA methylation analysis. DNA methylation levels were determined by bisulphite pyrosequencing. Briefly, 1 μg DNA were treated using the EpiTect Bisulfite Kit (Qiagen) and candidate-gene regions containing specific CpGs within *FKBP5* intron 7 (Paquette et al., 2014), *BDNF* untranslated exon IV (Perroud et al., 2013), and *NR3C1* exon 1F (Murgatroyd et al., 2015) were amplified using the PyroMark PCR Kit. See Table 1 for primer sequences, locations of regions, and PCR conditions. We focused only on specific CpGs supported by previous literature to maintain statistical power and reduce effects of multiple analyses. Single-stranded biotinylated product was purified by mixing 10 μl of the amplification mixture, 2 μl of streptavidin sepharose HP (Amersham Biosciences), and 40 μl of binding buffer. The sepharose beads containing the immobilized biotinylated product were purified, washed, and denatured in 0.2 mol/l NaOH and washed again using the Pyrosequencing Vacuum Prep Tool (Qiagen). The biotinylated DNA was resuspended in 12 μl of annealing buffer containing 0.3 μmol/l pyrosequencing primer (see Table 1 for primer sequences) and quantified by pyrosequencing using the PSQ 24MA system with the PyroMark Q24 Advanced CpG Reagents (Qiagen). The percentage methylation for each of the CpG sites was calculated using Pyro Q-CpG software (Qiagen). All analyses represent the average of three separate assays.

**Table 1.**
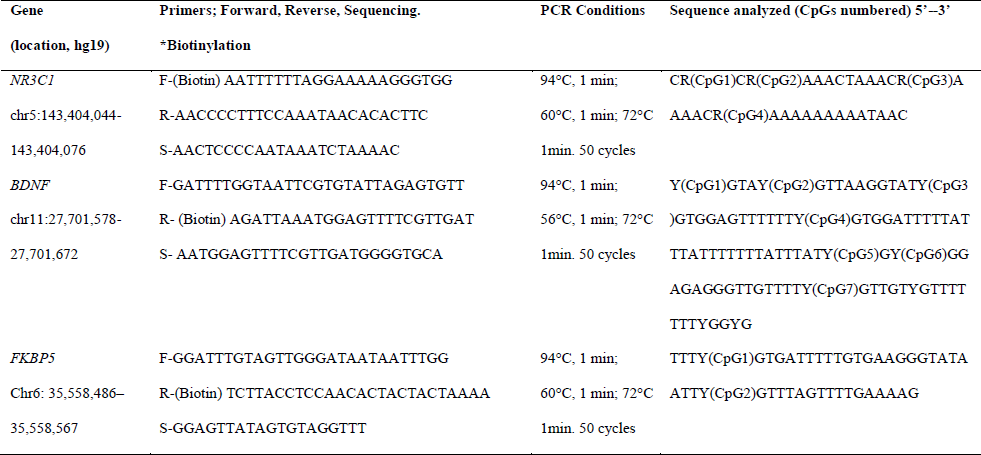
Primer sequences, locations and sequences of regions targeted by bisulphite pyrosequencing.

#### 2.2.3 Covariates

Maternal age, marital status, education, household income, ethnicity, years living in the US, nativity (US or non-US-born) and sex of the infant were collected through questionnaires at T1. We controlled for infant sex because previous literature suggests that newborn sex may be a key factor affecting the assimilation of prenatal stress into the epigenome (Braithwaite et al., 2015). These sex differences are likely to underlie the higher levels of glucocorticoids observed in females compared to males in response to acute and repeated stress (Seale et al., 2004). Because psychological distress is highly prevalent in Latina mothers and related to both discrimination and DNA methylation of stress-related genes (Berger and Sarnyai, 2015), we used the Inventory of Depression and Anxiety Symptoms – General Depression Scale (IDAS-GD) (Watson et al., 2012) which comprehensively assesses depressive symptoms to account for negative mood at T1 and T2; higher IDAS-GD scores indicates more severe symptoms. Typical IDAS-GD scores are 32.4 and 37.4 for control and high-risk women, respectively, and between 44.6 and 57.3 for depressed women (Schiller et al., 2013; Segre et al., 2015). Cronbach’s alpha for item consistency for the IDAS-GD in our sample was > 0.78 for T1 and T2.

#### 2.2.4 Statistical Analysis

We set out to model the associations between DNA methylation in the prenatal period and concurrent and later EDS scores (outcome) with the goal of understanding their relationship over time. We modelled the composite EDS scores with zero-inflated Poisson (ZIP) models (Lambert, 1992) because of overdispersion sourced from a high frequency of zero counts in this score. The ZIP model fit the data more consistently than other models considered (i.e., negative binomial and zero-inflated negative binomial models). ZIP regression was used to model count data that has an excess of zero counts and assumes that the excess zeros can be modelled separately from the count values. Specifically, the ZIP regression model has two parts, a Poisson regression models for the counts, and a logistic model for excess of zeroes. In our study, the logistic model has only one covariate, IDAS-GD, as it was the only covariate associated with the excess zero process. We examined the association between CpG methylation and EDS by including specific CpG’s in the Poisson model for counts, controlling for the variables of age, sex of baby, marital status, education, total income, ethnicity, years living in the US, and IDAS-GD score. We used the Vuong test (Vuong, 1989) to compare the ZIP with an ordinary Poisson regression model in terms of model fit. We used a post-hoc adjustment for multiple comparisons via the Benjamini-Hochberg procedure and controlled for the false discovery rate of 0.05. Only complete cases of covariates and control variables were considered (n = 147). Model goodness of fit was measured via McFadden pseudo-R^2^. McFadden pseudo-R^2^ measures the proportion of the variance in the outcome explained by the covariates, much like the coefficient of determination in an ordinary least squares (OLS) model. Instead of using sums of squared errors to construct R^2^, as in an OLS model, we used the log-likelihoods of the full and null models. We present Poisson model-based risk ratios associated with each CpG site and respective *p*-values, adjusted post-hoc via the Benjamini-Hochberg procedure, and McFadden pseudo-R^2^ values. To facilitate replication of this study, the R analytical code is available in Appendix A and the data can be downloaded from this link: https://osf.io/am58g/.

#### 2.2.5 Missing Data

A monotone missingness pattern was observed in follow-up (T2) IDAS-GD, and missing observations were multiply imputed with chained equations via predictive mean matching (White et al., 2011). Three observations with missing covariates and DNA methylation data for some of the markers were dropped from the study to avoid inducing bias due to imputation of both predictor and covariates.

## 3. Results

Table 2 summarizes the demographics of the cohort of 150 Latina women who were included in this analysis. The majority of our participants (78.8%) chose to complete data collection in Spanish. Participants had a mean age of 27.6 years. Most participants were married or living with a partner (74.2%), had an education level of high school or less (85.0%), and had a yearly household income of  25,000 US dollars (79.6%). The majority were non-US born (83.7%) and had been living in the US for a mean of 12 years. Of the sample, 56.3% were of Mexican origin, 17.2% were of Honduran origin, 13.3% were of Salvadoran origin, and the remaining 13.4% identified with other Latin American countries. In terms of depression symptoms from the IDAS-GD, the mean score was 29.87. In our sample, 43.5% of the women reported having experienced some type of discrimination (EDS > 0), with 30.4% of the women specifically reported experiencing ethnicity-based discrimination at some point in their lives. For the EDS outcome, the mean scores were 0.34 (SD 0.56) and 0.22 (SD 0.46) at T1 and T2, respectively. The most frequently reported reasons for experiencing discrimination were race and ancestry at both time points (Table 3).

**Table 2.**
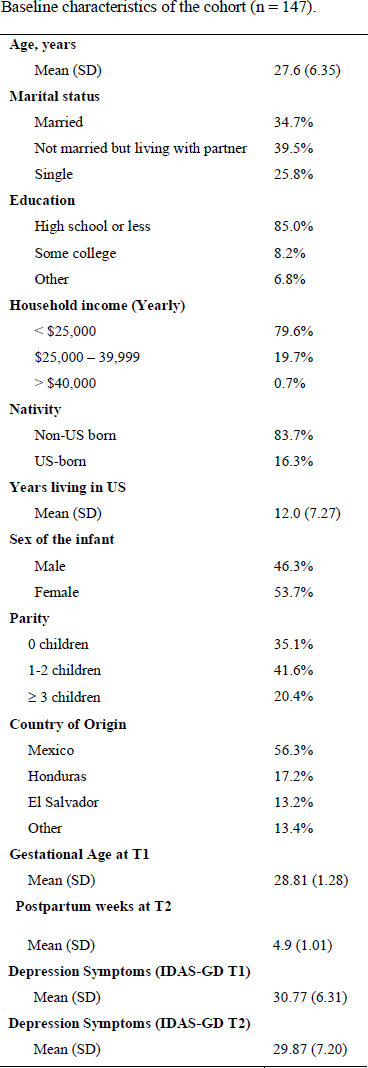
Baseline characteristics of the cohort (n = 147).

|  |  |
| --- | --- |
| **Age, years** |  |
| Mean (SD) | 27.6 (6.35) |
| **Marital status** |  |
| Married | 34.7% |
| Not married but living with partner | 39.5% |
| Single | 25.8% |
| **Education** |  |
| High school or less | 85.0% |
| Some college | 8.2% |
| Other | 6.8% |
| **Household income (Yearly)** |  |
| < \$25,000 | 79.6% |
| \$25,000 – 39,999 | 19.7% |
| > \$40,000 | 0.7% |
| **Nativity** |  |
| Non-US born | 83.7% |
| US-born | 16.3% |
| **Years living in US** |  |
| Mean (SD) | 12.0 (7.27) |
| **Sex of the infant** |  |
| Male | 46.3% |
| Female | 53.7% |
| **Parity** |  |
| 0 children | 35.1% |
| 1-2 children | 41.6% |
| ≥ 3 children | 20.4% |
| **Country of Origin** |  |
| Mexico | 56.3% |
| Honduras | 17.2% |
| El Salvador | 13.2% |
| Other | 13.4% |
| **Gestational Age at T1** |  |
| Mean (SD) | 28.81 (1.28) |
| **Postpartum weeks at T2** |  |
| Mean (SD) | 4.9 (1.01) |
| **Depression Symptoms (IDAS-GD T1)** |  |
| Mean (SD) | 30.77 (6.31) |
| **Depression Symptoms (IDAS-GD T2)** |  |
| Mean (SD) | 29.87 (7.20) |

**Table 3.**
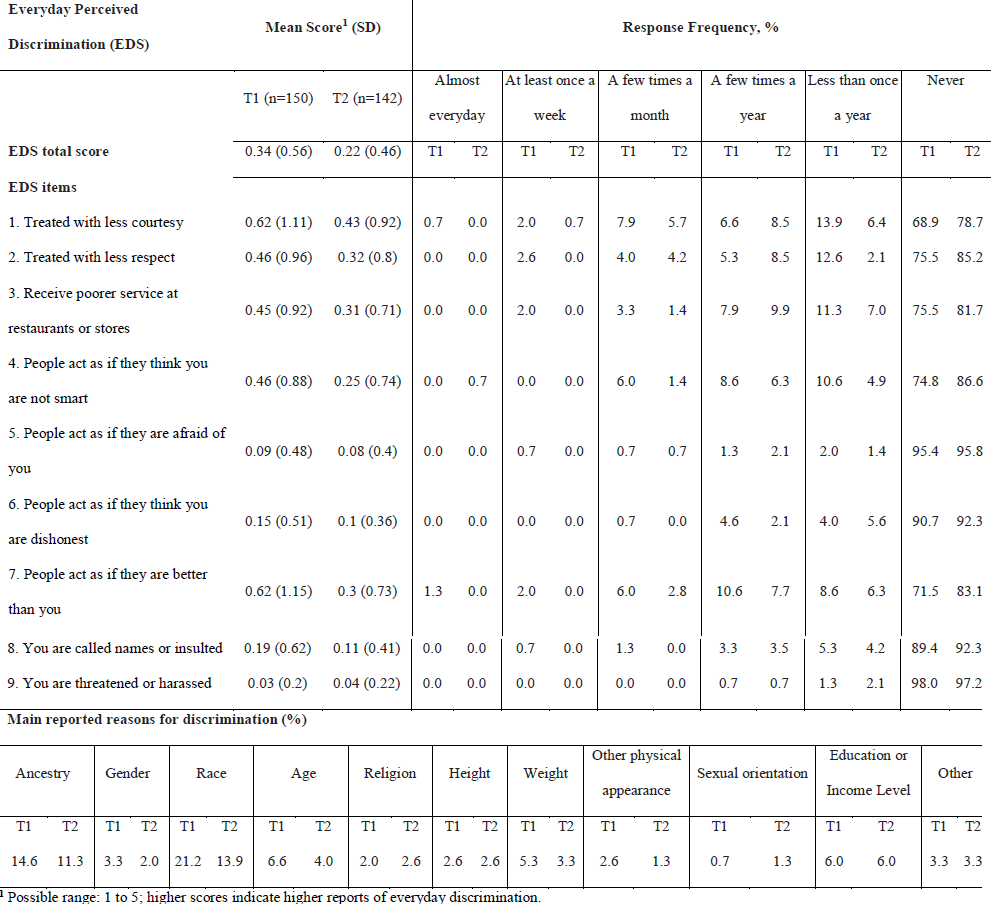
Response Frequencies and Mean Scores for Items on the Everyday Discrimination Scale (EDS).

Evidence of one-way association between EDS and methylation at various sites suggested further modeling with potentially confounding variables (Appendix B, Figure 1). Vuong tests to compare non-nested models indicated that ZIP is a better fit to our data than ordinary Poisson regression, all with *p* < 0.001. Table 4 shows the risk ratios between methylation at CpG site and EDS, estimated from the Poisson model part, controlling for all covariates listed previously (maternal age, marital status, education, household income, ethnicity, years living in the US, nativity, sex of the infant and mood symptoms). At T1, we found significant negative associations between EDS and methylation at CpG sites 1 and 2 of *NR3C1* (RR = 0.85, 0.84 and *p* = 0.008, 0.004, respectively). Significant negative associations were also identified at CpG sites 6 and 7 of the *BDNF* promoter (RR = 0.86, 0.92, *p* = 0.004, 0.004, respectively). Lastly, a significant negative association at CpG site 1 of *FKBP5* was identified (RR = 0.85, *p* < 0.001). At T1, significant covariates were sex of the baby (associated with decreased EDS risk), and absence of a partner, income greater than US$40,000, years living in the US, and depressive symptoms (associated with increased EDS risk).

**Table 4.** Estimated risk ratios between baseline (T1) and follow-up (T2) EDS and methylation at various CpG sites, controlled for demographic and mood symptom variables. P-values are from partial Wald-type added-last tests of regression coefficients and adjusted for multiple comparisons post-hoc via the Benjamini-Hochberg procedure, controlling for a 5% FDR. McFadden pseudo-R^2^ are provided for model fit.

| Gene | CpG Site | Genome location (hg38) | EDS at T1 |  |  | EDS at T2 |  |  |
| --- | --- | --- | --- | --- | --- | --- | --- | --- |
|  |  |  | Risk Ratio (95% CI) | p-value | Pseudo-R<sup>2</sup> | Risk Ratio (95% CI) | p-value | Pseudo-R<sup>2</sup> |
| **NR3C1** | Site 1 | chr5:143,404,073 | 0.85 (0.768,0.947) | **0.008** | 0.170 | 0.98 (0.826,1.154) | 0.891 | 0.145 |
|  | Site 2 | chr5:143,404,071 | 0.84 (0.752,0.931) | **0.004** | 0.174 | 0.84 (0.741,0.948) | **0.025** | 0.167 |
|  | Site 3 | chr5:143,404,061 | 0.92 (0.848,1.002) | 0.110 | 0.162 | 0.71 (0.471,1.066) | 0.234 | 0.187 |
|  | Site 4 | chr5:143,404,056 | 0.95 (0.867,1.034) | 0.308 | 0.158 | 0.93 (0.794,1.080) | 0.442 | 0.148 |
| **BDNF** | Site 1 | chr11:27,701,672 | 1.03 (0.982,1.073) | 0.308 | 0.158 | 1.00 (0.886,1.140) | 0.937 | 0.144 |
|  | Site 2 | chr11:27,701,668 | 1.02 (0.974,1.063) | 0.477 | 0.156 | 1.05 (0.988,1.118) | 0.234 | 0.151 |
|  | Site 3 | chr11:27,701,657 | 0.91 (0.822,1.008) | 0.123 | 0.161 | 0.88 (0.744,1.035) | 0.234 | 0.151 |
|  | Site 4 | chr11:27,701,644 | 0.96 (0.931,0.996) | 0.065 | 0.163 | 0.98 (0.938,1.026) | 0.503 | 0.147 |
|  | Site 5 | chr11:27,701,616 | 0.96 (0.882,1.042) | 0.376 | 0.157 | 0.86 (0.763,0.958) | **0.025** | 0.161 |
|  | Site 6 | chr11:27,701,612 | 0.86 (0.793,0.939) | **0.004** | 0.176 | 0.89 (0.819,0.962) | **0.025** | 0.167 |
|  | Site 7 | chr11:27,701,597 | 0.92 (0.880,0.967) | **0.004** | 0.175 | 0.92 (0.880,0.971) | **0.025** | 0.169 |
| **FKBP5** | Site 1 | chr6:35,558,566 | 0.85 (0.787,0.919) | **< 0.001** | 0.180 | 0.97 (0.907,1.031) | 0.442 | 0.147 |
|  | Site 2 | chr6:35,558,514 | 0.98 (0.945,1.012) | 0.305 | 0.158 | 0.96 (0.893,1.025) | 0.354 | 0.148 |

At T2, consistent with the findings at T1, the negative associations between EDS and CpG site 2 of *NR3C1* (RR = 0.84, *p* = 0.025) and CpG sites 6 and 7 of *BDNF* (RR = 0.89, 0.92, and *p* = 0.025, 0.025, respectively) were still present. The analysis also showed negative associations of EDS with CpG site 5 of *BDNF* (RR = 0.86, *p* = 0.025). At T2 significant covariates were age and presence of a partner (associated with decreased EDS risk), and income and general depression (associated with increased EDS risk). The complete model results from T1 and T2 are presented in Appendix B, tables B1-B4.

Considering the present results, we tested whether ethnicity-based discrimination was associated with DNA methylation for each CpG site in a post-hoc analysis. However, a basic test of association via logistic regression showed no association between average DNA methylation at these sites with perceived ethnic discrimination (Appendix B, Figure 2).

## 4. Discussion

This is the first study to report associations between blood DNA methylation of stress-related genes (*NR3C1, FKBP5, BDNF*) and perceived discrimination in Latina women in the US. Exposure to discrimination has established adverse impacts on health. We hypothesized that DNA methylation within the *NR3C1, FKBP5*, and *BDNF* genes would be inversely associated with perceived discrimination. In our cohort, 43.5% of the women reported having experienced discrimination of some sort. Via the EDS, women reported low to moderate frequency of discriminatory experience, which is consistent with a previous study (Colen et al., 2018), and related their discrimination experiences mostly to their race and ancestry. We identified several statistically significant associations, even after accounting for a stringent list of covariates, including demographics, ethnicity, immigration and mood symptoms. Our findings underscore the specific and complex biological pathway associations between discrimination and epigenetic modification in Latina women in the US.

Within the *NR3C1* exon 1F, methylation at CpG site 2 was negatively associated with EDS at both T1 and T2 while CpG 1 methylation was negatively associated with EDS at T1. The *NR3C1* exon 1F is a key element in stress response regulation, and the present data suggest that increased EDS is associated with regulatory changes in glucocorticoid-related genes. The fewer associations at T2 are likely to be due to the 18-week gap between methylation levels at T1 and EDS assessment at T2. Other potential explanations include the endocrine changes during the peripartum period (including elevated cortisol levels) and/or changes in EDS perception due to motherhood and associated improvements in the social environment (decreased exposure to negative social interactions, including discrimination, and increased exposure to social support). These explanations need to be explored in future studies. Our finding that some of the associations between DNA methylation and EDS score hold over time deserves further investigation as DNA methylation markers identified could serve as a risk factor and/or biomarker for mothers at risk of the adverse effects of discrimination.

Differences in methylation patterns in the *NR3C1* exon 1F (or Exon 1_7_ in rats) in relation to social environment and stress have been reported in a systematic review of 40 articles (27 human and 13 animal studies) (Turecki and Meaney, 2016). In studies focused on psychological distress, human studies (seven in total) reported varied results in terms of *NR3C1* exon 1_7_ methylation: one reported increased methylation (Dammann et al., 2011), two reported decreased methylation (Alt et al., 2010; Yehuda et al., 2015), while three reported no change (Alt et al., 2010; Steiger et al., 2013; Yehuda et al., 2013). Similar findings were also reported in early life stress and parental stress studies. For example, in a socioeconomic-matched analysis of children exposed to maltreatment, researchers found decreased methylation in a single CpG (CpG 2, corresponding to our CpG 1) and increased methylation at CpGs 3, 5 and 6 (Romens et al., 2015). Another study found that maternal and paternal experience of the Holocaust were associated with decreased and increased methylation of exon 1F, respectively (Yehuda et al., 2014). A meta-analysis further demonstrated changes at specific CpG 36 (corresponding to our CpG 1) site and prenatal stress in infants supporting the importance of methylation at key CpG sites within *NR3C1* (Palma-Gudiel et al., 2015). We hypothesize that Latinas that perceive higher levels of discrimination may not effectively down regulate glucocorticoid levels in response to this type of stress, resulting in increased susceptibility to adverse health outcomes. This discrimination exposure could be mediated by both genetic susceptibility and/or exposure to chronic stress. The current *NR3C1* methylation results indicate that changes in glucocorticoid methylation may be a potent risk factor and support the investigation of this specific gene.

Another important modulator of glucocorticoid signaling in response to stress is *FKBP5*, and the data suggest that the methylation of this glucocorticoid binding protein as well as the receptor gene are inversely associated with elevated EDS. The *FKBP5* and *NR3C1* data together indicate that HPA related methylation changes are associated with both pre and postpartum discrimination, but the specific nature of this relationship may vary with time. The EDS and methylation findings support the hypothesis of increased responsiveness to social stress in subjects with higher EDS scores. One speculative implication of these data is that discrimination-related stress could induce coordinated DNA methylation effects on multiple genes that collectively serve to downregulate stress responsivity. However, a key consideration in these types of studies is that those who volunteer may tend to be relatively more resilient, less sensitive to discrimination, and express fewer psychological distress. While the design of the present study did not allow for causal mechanistic analyses, future studies should investigate the DNA methylation regulation of these stress-related genes in a larger and more heterogenous cohort with a prospective design.

Significant negative associations between EDS and *BDNF* methylation were also observed. BDNF is a major mediator of neuronal plasticity and there is substantial evidence that *BDNF* expression and neurogenesis are generally reduced following chronic and acute stressors in human and animal studies. The negative association at T1 and T2 (CpGs 6 and 7) and potential increase in *BDNF* in those exposed to higher levels of EDS may be driven by the type of BDNF actions associated with post-traumatic stress (Zhang et al., 2016), where increases may consolidate the behavioral effects of adverse stressful events through neuroplasticity mechanisms. It is interesting to note that behavioral and neural changes observed in mothers who experience a traumatic birth are similar to those found in patients with post-traumatic stress disorder (Yildiz et al., 2017).

A further supposition is whether discrimination might be positively adaptive in the sense of heightening awareness and attention to the environment. Such components of consciousness have been suggested to influence brain neuroplasticity, activating synaptic flow and changing brain structures and functional organization (Askenasy and Lehmann, 2013). The present data and the BDNF literature further underscore the imminent need for long-term prospective studies of the role of BDNF in the etiology of stress-related disorders in Latinas exposed to discrimination.

Looking at the covariates included in the regression models, we observed that sex of the baby (T1), age and presence of a partner (T2) all decreased the risk ratios for EDS, while depressive symptoms and higher annual income (T1 and T2), years living in the US and being single (T1) increasing the risk of discrimination. These findings reinforce reports of several risk factors for stress-related disorders and indicate that increased age and partner support may be particularly and specifically protective against the risk of discrimination in the perinatal period. Furthermore, increased risk associated with years living in the US may be due to impaired socio-cultural based resilience in these individuals (Cardoso and Thompson, 2010). The strikingly substantial increased risk of discrimination associated with greater income level extends the findings of previous studies showing an association between greater discrimination and increases in income (Colen et al., 2018). This suggests that potential benefits of the socioeconomic status gradient derived from greater income and education status may not be uniformly protective against social stressors, such as discrimination. Given the current methylation data, the association between depressive symptoms and EDS risk, and related research on the potency of discrimination as a social stressor, future studies should explore whether the relationship between discrimination and DNA methylation is mediated by HPA related factors and mechanisms. There is also a
need to carefully consider social and temporal based characteristics of study populations to explore their potential effects between stress-related DNA methylation and discrimination.

Given the evidence of the role of elevated cortisol levels in the adverse effects of discrimination on mental health (Berger and Sarnyai, 2015; Zeiders et al., 2012), the elevated prevalence of perinatal depression in Latinas (Gentile, 2017; Liu and Tronick, 2014), and the adverse effects of peripartum cortisol (Bergman et al., 2010), other stress related factors (Beijers et al., 2014) on offspring development and health, it is highly probable that discrimination can have negative effects on both mother and child. It is also possible that discrimination, as a robust and prevalent social stressor, may be a primary contributing factor in the high rates of Latina psychological distress (Halbreich and Karkun, 2006; Liu and Tronick, 2014). Assessment of discrimination in parents may serve as a sensitive and highly relevant indicator of elevated risk for perinatal depression and anxiety and the associated negative consequences on offspring.

Some limitations need to be taken into consideration while interpreting the results of this study. First, we focused on key CpGs to limit the potential impact of multiple analyses. However, there are many other CpG sites and combinations that could be explored. Second, we analyzed methylation within peripheral blood samples. We must consider that blood is heterogeneous, which may account for some of the variability in methylation and may introduce a confound where other variables are associated with cellular heterogeneity. Third, while studies combining methylation in blood and post mortem brain suggest that they are often substantially correlated (Tylee et al., 2013), it cannot be assumed that DNA methylation in peripheral tissues reflects methylation in relevant central nervous system regions. This is particularly a concern due to substantial variation in epigenetic effects across brain regions and cell types. Fourth, we used a self-reported measure of discrimination, thus introducing the risk of report bias. The EDS assesses discrimination across several domains, without specific reference to race, ethnicity or other demographic characteristics. This feature of the EDS allows it to be used across populations of different racial/ethnic backgrounds and also allows us to tap into the subjective experience of perceived discrimination (Lewis et al., 2012). While the overall sample size of 147 is modest, larger studies are needed to explore epigenome-wide analysis. Our data collection was completed between May 2016 to March 2017, which overlaps with the 2016 U.S. presidential election in which Latin American immigration to the US emerged as one of the most politicized and polemical topics on the campaign trail. Thus, it is possible that the reports obtained from our assessments, especially at T2 (post-election), were affected by the increased self-awareness and self-protection of Latinos within our communities, which could explain the small decrease in EDS report from T1 to T2. Answering reports based on social desirability prevents participants from increasing their interaction with the research team and related health care providers (Hopwood et al., 2009). In this sense, the EDS scores reported in this report are likely an underestimation of the actual experience of this population. Future studies should consider using multiple data collection methods to capture the complex nature of discrimination in an individual’s life and social desirability on their approach to self-report social and health-related information.

In summary, our findings indicate that discrimination exposure is inversely associated with DNA methylation intricately involved in the etiology of stress-related disorders, such as depression, anxiety, and post-traumatic stress, which substantially and disproportionally affect Latino communities. There were differences in methylation patterns within and across genes, emphasizing the importance of specificity in methylation patterns among CpG sites and reinforcing the call for studies to target CpG sites within biologically relevant areas, such as transcription factor binding regions and non-coding first exons of the *NR3C1* gene. In addition, there is a need for expression studies to determine the functional repercussions of CpG methylation. These results warrant further investigation to better understand the genetic and psychopathological impact of discrimination on Latino mothers and their families.

## Contributors

Authors HS, BN and CM designed the study. Authors HS, BN, LS, RA, and CM were involved in data collection, processing and/or quality assurance. Authors AB, XT, and HS performed the statistical analyses and/or made tables. All authors contributed to the interpretation of the results. Author HS wrote the first draft of the manuscript. All authors contributed to and have approved the final manuscript.

## Acknowledgements

We would to thank our research assistants, Erika Campos and Kathia Pena, for their effort in the recruitment and retention of participants in this study, and Victoria Benson for her effort in helping manage our samples in the UNC Biobehavioral Laboratory.

